# The looping lullaby: closed-loop neurostimulation decreases sleepers’ sensitivity to environmental noise

**DOI:** 10.1101/2021.08.07.455505

**Authors:** Vidushi Pathak, Elsa Juan, Reina van der Goot, Lucia M. Talamini

## Abstract

**Study Objective:** Sleep is critical for physical and mental health. However, sleep disruption due to noise is a growing problem, causing long-lasting distress and fragilizing entire populations mentally and physically. Here for the first time, we tested an innovative and non-invasive potential countermeasure for sleep disruptions due to noise.

**Methods:** We developed a new, modeling-based, closed-loop acoustic neurostimulation procedure (CLNS) to precisely phase-lock stimuli to slow oscillations (SO). We used CLNS to align, soft sound pulses to the start of the SO positive deflection to boost SO and sleep spindles during non-rapid eye movement (NREM) sleep. Participants underwent three overnight EEG recordings. The first night served to determine each participant’s individual noise arousal threshold. The remaining two nights occurred in counterbalanced order: in the “Disturbing night”, loud, real-life noises were repeatedly presented; in the “Intervention night”, similar loud noises were played while using the CLNS to boost SO. All experimental manipulations were performed in the first three hours of sleep; participants slept undisturbed for the rest of the night.

**Results:** In contrast to the Disturbing night, the probability of arousals caused by noise was significantly decreased in the Intervention night. Moreover, the CLNS intervention increased NREM duration and sleep spindle power across the night.

**Conclusions:** These results show that our CLNS procedure can effectively protect sleep from disruptions caused by noise. Remarkably, even in the presence of loud environmental noise, CLNS’ soft and precisely timed sound pulses played a beneficial role in protecting sleep continuity. This represents the first successful attempt at using CLNS in a noisy environment.

**Statement of Significance:** Exposure to noise during sleep impairs sleep quality, leading to impaired biological functioning, decreased day-time performance and higher occurrence of accidents. Slow wave sleep is hallmarked by slow oscillations (SO) and sleep spindles, both of which are markers of sleep stability. In previous experiments, acoustic stimulation has shown potential in enhancing SO and sleep spindles. Whether these manipulations also work to stabilize sleep against the disrupting effects of environmental noise remains unclear. Here we use a new closed-loop approach to precisely align subtle acoustic stimuli with SO phase. We show that this method effectively stabilizes sleep in the presence of noise. Our results bear crucial relevance for improving sleep in the general public and in many high-risk professions.

## 1. Introduction

Good sleep quality is essential for physical and mental health. Deep sleep or slow wave sleep (SWS), in particular, plays an important role in many biological processes^1, 2^ supporting growth, restoring the immune system and regulating inflammation ^1 2^.

Accordingly, sleep deficiency can over time, increase the risk of diabetes, cardiovascular disorders and psychiatric disorders. Sleep deprivation also leads to daytime sleepiness, impaired motor skills and cognitive and emotional deficits^3^, which drastically increase the risk of human errors and cause about 20% of road accidents ^4^ and 36% of medical errors ^5, 6^.

The above is particularly relevant in this day and age, considering that adults in industrialized areas across the world are operating at a massive sleep deficit: a north American adult typically averages a total of about 6 hours of sleep a night ^4^, while average sleep need amounts to approximately 7-9 hours ^7^.

One of the most common causes of sleep disturbance is the presence of noise in the sleeper’s environment. Indeed, exposure to noise during sleep increases the number of arousals and decreases time in SWS, leading to poor sleep quality ^8–10^ and increased sleepiness during the day ^10^. Unfortunately, environmental noise is ubiquitous in modern societies, especially in city centers, near construction sites, airports and train stations, but also in hospital rooms, with adverse consequences for patient recovery ^9^. Sleep quality tends to be particularly poor in ICU environments, and noise seems to be among the top causes ^11^. Finally, sleep fragmentation due to noise is prevalent in many high-demand jobs, such as in the military or on spaceflight missions ^12^, leading to poor cognitive performance ^13, 14^ with potentially tremendous consequences. Since it is not always possible to avoid environmental noise, it is imperative to find solutions to reduce its impact on sleep.

Slow oscillations (SO; 0.5-1 Hz) are a hallmark of non-rapid eye movement (NREM) sleep and a robust indicator of sleep depth: deeper the sleep, higher the arousal threshold to external sensory stimuli. The underlying mechanism involves a decreased thalamic responsiveness to sensory signals, which increases as sleep deepens, reducing cortical responsiveness to such stimuli^15^. This gating of afferent signals is particularly effective when sleep spindles (11-16 Hz) are present^16^. Accordingly, spindles have been shown to play a role in resistance to external sounds^17^. Interestingly, both SO and sleep spindles can be enhanced - at least momentarily - by external stimuli, including rocking^18^, transcranial magnetic^19^ or alternating current stimulation^20^ and acoustic stimulation^21–23^. Such findings have inspired the notion that sleep might be improved through external stimulation procedures. The use of acoustic stimuli present advantages pertaining to safety, ease of use and miniaturization, that are particularly promising for the development of clinical and home-use therapies^22–27^.

A particularly effective way to stimulate SO was developed in our lab. The procedure, termed closed-loop neurostimulation (CLNS), involves flexible, near real-time modeling of the EEG signal’s oscillatory dynamics ^22, 28^. This allows prediction of upcoming brain activity and precise targeting of features of interest, such as oscillatory phase. Recent studies from our lab have shown that subtle, non-arousing sounds targeted at the start of the positive SO halfwave (hereafter referred to as 0° phase) can boost the endogenous SO dynamic, increasing the amplitude of the following SO, inducing long trains of SO and extending deep sleep ^45^. Using CLNS, we have shown that 0° phase targeting also maximizes the induction of sound-evoked spindle activity, compared to a control phase manipulation ^22^.

Here we aim to investigate whether CLNS could counteract sleep disturbances due to noise. We used CLNS in sleep stages N2 and N3 while loud environmental noises were regularly presented. In a within-subjects design, we recorded high-density EEG on three non-consecutive nights: (1) a ‘threshold’ night to determine subjects’ individual noise arousal threshold, (2) a night with repeated presentation of loud noises (Disturbing night) and (3) a night where, in addition to noise, CLNS was used (Intervention night).

As an individual’s arousal threshold is highly dependent on the depth of sleep, disturbing noises were only presented during deep sleep (stage N3). The sleep disruption protocol was interactive, with noises being presented as long as the subject remained in deep sleep and halted as soon as the subject left deep sleep, be it due to an arousal or a transition to another sleep stage. Therefore, any CLNS-induced sleep stabilization could be reflected both in fewer arousals and in more noise stimuli being presented.

In accordance with the above, the main outcome measure of this study was the arousal probability (ratio of arousals to noises). We hypothesized that CLNS would lead to a decrease in the probability of arousals due to noise. The main outcome measure of this study was the arousal probability (ratio of arousals to noises). We hypothesized that CLNS would lead to a decrease in the probability of arousals due to noise. While the experimental approach did not capitalize on assessing the effects of CLNS intervention on sleep measures (because the number of disturbing noises was not equal during the two experimental nights), a number of such measures was nevertheless assessed. In particular, we analysed sleep architecture, as well as frequency power measures during NREM sleep, to assess whether CLNS would boost SO and sleep spindles and increase sleep continuity in the Intervention night. Measures of subjective sleep quality were also assessed.

## 2. Methods

### 2.1 Participants

Twelve healthy participants participated in this study. Two participants for whom the CLNS procedure was sub-optimal (< 300 boosting sound pulses delivered) were removed such that 10 healthy participants (median age [21] and range [18 - 30]; 7 females) were included in the final analysis. All participants were free of medical problems, subjective sleep issues, and were selected for reporting habitual good sleep quality. An audiometric screening was done to confirm normal hearing. Participants were instructed to keep a regular sleep window between 23:00 and 08:00 for at least one day before coming to the lab, not nap in the daytime, and to wake up no later than 08:00 on the morning of each experimental night. The interval between experimental nights was between 2 and 8 days.

To keep participants naïve of the study’s expectations, they were informed that the goal of the experiment was to study real-world sound processing during sleep. Thus, they knew that sounds would be presented during the night, but not the nature of the sounds, nor that some sounds might boost their sleep. This study was conducted according to the principles of the Declaration of Helsinki. Procedures were approved by the local ethics committee and all participants provided written informed consent. Participants were compensated with credits to fulfil course requirements or money.

### 2.2 Stimuli

Ten sleep disturbing sounds of 10 s each, representing real-world noises ("good" conversation, "bad" conversation, paging, door opening and closing, telephone ringing, laundry cart rolling, helicopter take-off, jet engine flyover, IV pump alarm and traffic flow) were selected from Buxton et al. (2012) ^9^. These sounds were modified using Audacity software (http://audacityteam.org) to include a fading in and fading out effect of 10 ms each.

CLNS used a 5 ms ‘sound pulse’, played at an unobtrusive volume of 42 dB. Two speakers, placed 50 cm from the subject’s head, were used to play both the noise stimuli and the CLNS sound pulses that sound like ‘clicks’.

### 2.3 Experimental Design and Procedure

On each experimental night, participants arrived at the sleep laboratory at 21:30 and were hooked up for standard high-density polysomnography (see 2.6 Data acquisition). Lights were turned off at 23:00 and participants were allowed to sleep until 09:00 the next day. Real-time EEG signals were constantly monitored by the experimenter. On all three nights, noise presentation occurred in N3 during the first two sleep cycles (∼3h).

The first night was always a threshold night, aimed at determining each participant’s arousal threshold (see Figure 1 for a schematic overview of the experimental procedures). Two types of noises (‘traffic’ and ‘good conversation’) were played alternatingly. Volumes ranged between 40 dB and 70 dB with increments of 5 dB and an interstimulus interval of 10 s. Arousals during the stimulation period were scored off-line according to AASM guidelines^29^. A participant’s arousal threshold was determined as the minimal volume eliciting an arousal at least 50% of the time. This individual arousal threshold was used to establish the volumes at which the noise stimuli would be presented during the two subsequent experimental nights.

**Figure 1.**
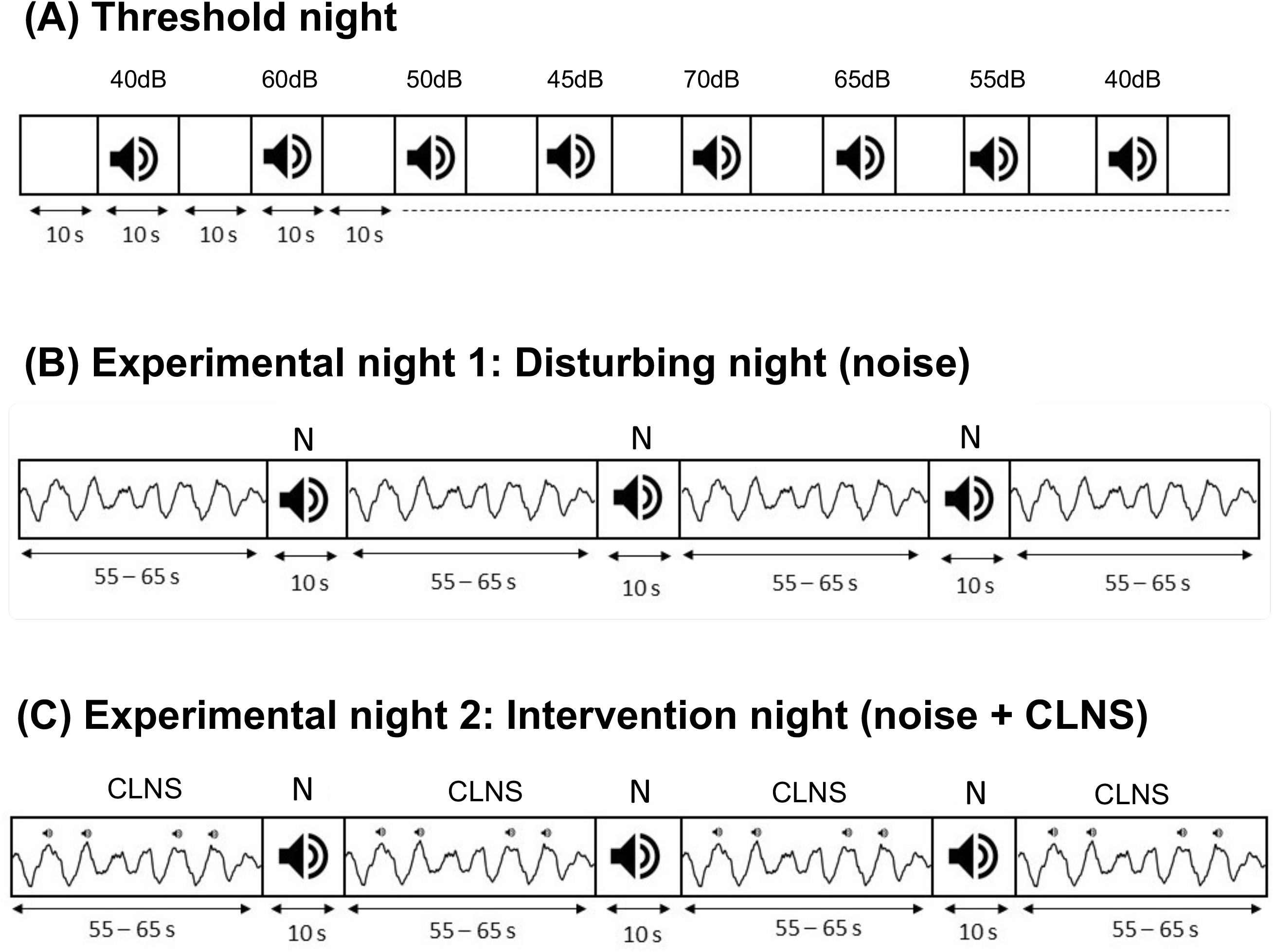
Schematic illustration of the stimulation period for the three laboratory nights. (A) The threshold night was the first night for all participants. Individual arousal thresholds were measured by presenting noises of 10 s duration (N) at various sound intensities during N3. Noises were separated by 10 s inter-stimulus intervals to assess possible arousals or awakenings. (B) The order of two following experimental nights was counterbalanced across participants. In the Disturbing night, noises were played in stage N3 at a volume ranging ± 5dB around the participants’ individual arousal threshold with an inter-stimulus interval of 60s ± 5s. (C) In the intervention night, the same procedure was applied, but sound pulses phase-locked to the 0° phase of SO were presented throughout N2 and in the 60 ± 5 s. intervals between noises in N3. All experimental manipulations were performed in the first two cycles of sleep (‘stimulation period’, lasting ∼3 h) after which participants were able to sleep undisturbed for 6.5 h.

The order of the Disturbing night and Intervention night was counterbalanced across participants. To avoid habituation effects, each night used a different set of 5 disturbing sounds (set to condition matching was counterbalanced across participants). Noises in the two sets were matched on acoustic characteristics, meaning and potential to arouse, based on findings from Buxton et al. (2012), and pilot studies in our lab. During both nights, the first three sounds were presented 10 dB below the individualized arousal threshold, to avoid startling the participant at the beginning of the procedure. Thereafter, all noises were played at volumes ranging ± 5dB around the participant’s arousal threshold. The interval between noise stimuli was 60 ± 5s, randomly jittered. Noise presentation was paused whenever polysomnographic signs of N2, REM sleep, arousals or awakenings were observed; it was resumed upon renewed detection of N3.

During the Intervention night, in addition to the noise presentations, CLNS to boost sleep was applied during the first two sleep cycles, throughout N2 and N3 sleep. CLNS’ sound pulses were only paused during the presentation of noise stimuli.

### 2.4 CLNS procedure

In the Intervention night, we used our patented CLNS algorithm to target soft sound pulses at 0° phase (see Figure 2A) (adapted from Cox et al., 2014). A copy of the Fpz-M1 signal was transferred from the recording computer to the stimulation computer for real-time analysis. The data was received into a buffer holding the most recent 10 s of data. The CLNS algorithm then performed a non-linear sine fitting procedure on the most recent 1 s of unfiltered EEG signal and checked the following criteria for stimulus release at each iteration: (1) the fitted sine was in a predefined frequency range of interest (0.5-1.5 Hz), (2) the fitting error was below threshold of 0.3, (3) the predicted target phase occurred within a narrow time between 24 to 34 ms into the future. If all criteria were met, the sound pulse was released to coincide with the upcoming predicted target phase. To maximize CLNS boosting potential there were no imposed interstimulus intervals between the sound pulses. The real-time phase prediction algorithm was implemented in EventIDE software (Okazolab Ltd, London, United Kingdom).

**Figure 2.**
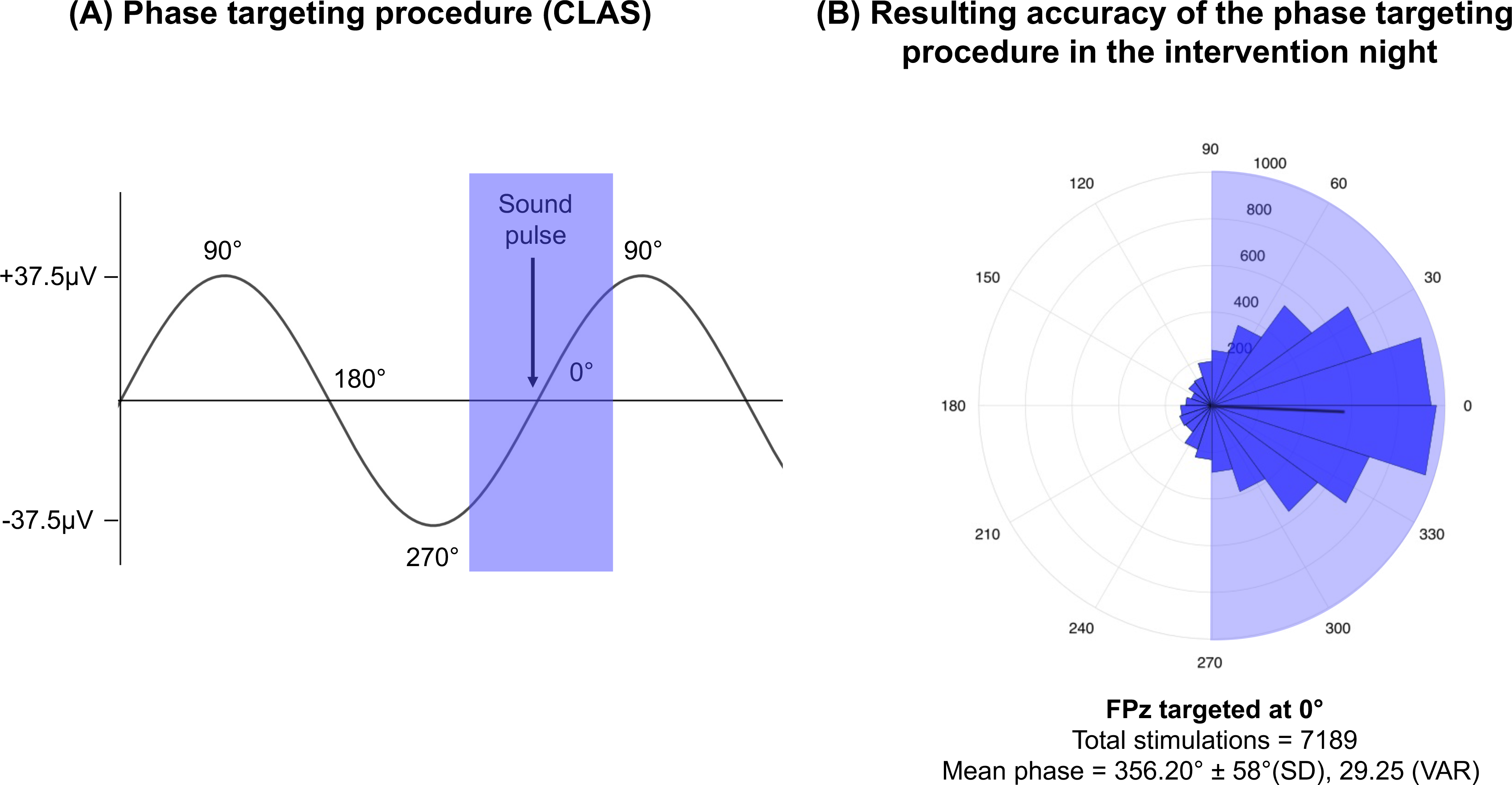
Phase targeting procedure and accuracy. (A) Our phase targeting procedure detected individual slow oscillations on the Fpz channel and delivered a soft, short sound pulse at the 0° phase of the incoming slow oscillation (start of the positive half-wave). (B) In total, this procedure resulted in 7189 stimulations across all participants (n=10). Phase at sound onset deviated only 3.8° from the 0° targeted phase, demonstrating high phase targeting precision.

### 2.5 Sleep Questionnaires

Participants’ subjective sleep experiences were assessed using three questionnaires: the Sleep quality questionnaire^30^, the Stanford sleepiness scale^31^ and the Consensus sleep diary^32^. All questionnaires were administered in the morning after each experimental night. The sleep quality questionnaire assesses the quality of sleep in the preceding night with 14 questions on a forced-choice scale. The Stanford sleepiness scale was administered to measure the level of sleepiness, ranging from 1 to 7. The consensus sleep diary has nine open-ended questions in total, with two main outcomes: the subjects’ perceived sleep quality and the perceived number of awakenings.

### 2.6 Data acquisition

High-density EEG was acquired using 62 electrodes placed according to the international 10-10 system (64-channels WaveGuard original cap, ANT, Enschede, The Netherlands) with two additional electrodes on the mastoids as reference. In addition, horizontal eye movements (HEOG), vertical eye movements (VEOG) and submental muscle tone (EMG) were measured with bipolar electrodes. Signals were sampled at 512 Hz using 72-channel Refa DC amplifiers and an in-house adapted version of Polybench software (TMSi, Oldenzaal, The Netherlands).

### 2.7 Data analysis

#### Sleep architecture

EEG data was re-referenced to the mastoids, high-pass filtered at 0.3 Hz and low-pass filtered at 35 Hz. Sleep stages were scored offline according to AASM criteria ^33^ using Galaxy software (PHIi, Amsterdam).

For each experimental night, we determined the duration (min) and proportion (%) of sleep stages N1, N2, N3, total NREM and REM sleep, as well as sleep efficiency, sleep latency, REM latency and time awake after sleep onset.

#### Arousals

In line with AASM criteria^33^, arousals during the stimulation period were defined as an abrupt shift of EEG frequency (e.g., alpha, theta) lasting at least 3 s, with at least 10 s of stable sleep preceding the change^30^. This includes movements, awakenings and shifts to a lighter sleep stage (N1/ N2). For this analysis, we only considered arousals related to noise, i.e., occurring within 10 s after noise presentation. For each subject, the total number of arousals across these combined time windows was assessed. In addition, a weighted arousal score was calculated using a point system to take into account the severity of arousals, such that bigger changes in sleep architecture were attributed a higher score. A score of 0.5 was attributed to movements during N2 and N3 that were not accompanied by a sleep stage shift. A shift in sleep stage from N3 to N2 was given a score of 1, N3 to N1 was given a score of 1.5 and awakenings from N3 were attributed a score of 2. Finally, for each participant, we calculated the probability of arousals due to noise as the weighted arousal score divided by the number of noises presented.

#### Preprocessing

Preprocessing of EEG data was done using MATLAB software (MATLAB 2016a, The Mathworks, RRID: SCR_001622) running custom scripts based on functions from the EEGLAB toolbox (EEGLAB 13.5.4, http://sccn.ucsd.edu/eeglab). The signal was high-pass (0.1 Hz) and low-pass (49 Hz) filtered, re-referenced to the average of the mastoids, down-sampled to 256 Hz and split in 30 s epochs. Power spectrum values were calculated for all 62 scalp electrodes on frequency bins of 0.25 Hz, ranging from 0.5 Hz to 48 Hz. Automatic trial rejection was performed using these values. To remove outliers from the data set, epochs with an average z-score larger than 2 were discarded. Further, epochs marked as containing arousals and muscle movements based on the AASM criteria^29^ were also discarded. As a final check, the signal was visually inspected; noisy channels were interpolated and artifactual epochs were rejected.

#### Differences in power across frequency bands

To examine the power changes induced by CLNS intervention, differences in power across frequency bands were computed on the NREM sleep data from electrode Fz during the whole night and in the stimulation period. The frequency content of the pre-processed EEG signal was decomposed in frequency bands using fast Fourier transform with Hamming window of 4 s, 50% overlap and 0.25 Hz bin size. The power per 30 s epoch was computed for each frequency bin and summated over all epochs.

Finally, relative power was calculated by dividing the power per frequency bin by the total power in the 0.5-48 Hz range. Relative power bins were merged across eight consecutive frequency ranges of interest: SO (0.5–1.5Hz), high-delta (1.5-4Hz), theta (4– 8Hz), alpha (8–11Hz), sigma (11–16Hz), beta (16–30Hz) and gamma (30–48Hz).

#### CLNS accuracy

To determine the precision of the CLNS phase-targeting procedure, we calculated the accuracy of sound pulses delivery in relation to SO phase. The signal of channel Fpz (used for stimulus targeting) was band-pass filtered between 0.5-1.5 Hz using a zero-phase shift Butterworth filter and a Hilbert transform was executed to extract the phase at each time point. The CircStat toolbox ^34^ was used to analyse the phase distribution of all delivered sound pulses.

#### Statistics

Differences in arousal measures, sleep architecture, power across frequency bands, and questionnaires were analyzed using paired samples *t*-tests (two-tailed) contrasting the Disturbing night and the Intervention night. Students’ *t*-test was used provided that assumptions were respected, otherwise Wilcoxon’s test was used.

## 3. Results

### 3.1 CLNS performance

On average, the CLNS procedure delivered 733 ± 113 sound pulses per participant, or about 6 ± 1 sound pulses per minute (Figure 2B). Taken together, this amounts to a total of 7189 sound pulses across the 10 participants during the 3h stimulation period in the Intervention night. The mean, standard deviation and variance of the slow wave phase at sound onset on Fpz was M = 356.20°, SD = 58°and VAR = 29.25° (Figure 2B). This means that the sound pulses were delivered on average at 4° from the aimed 0°, which demonstrates an extremely high phase-targeting accuracy.

### 3.2 Noise delivery

The number of delivered noise was 24 % higher in the Intervention night (42 ± 5.3) than in the Disturbing night (34 ± 4.8; *t* (9) = 1.48, *p* = 0.17). Though not statistically significant, this increase seems to suggest that CLNS procedure allowed the presentation of more noises. Indeed, if CLNS increases resistance to external sounds, lessens N3 interruptions and perhaps also speeds up return to N3 following arousals, then the probability of presenting noises becomes higher.

### 3.3 Arousals

On average, participants’ arousal threshold - as determined in the threshold night - was 50 ± 4 dB. To assess whether there was a decrease in arousals due to CLNS, we computed the sum and the probability of arousals due to noises in the Disturbing night and the Intervention night. On average, the CLNS intervention reduced the weighted arousal score by ∼51%, decreasing by half the number of arousals in the Intervention night as compared to the Disturbing night (4.95 ± 0.92 and 9.7 ± 0.89 respectively, *t* (9) = -3.98, *p* = .003) (Figure 3). The probability of arousals to occur due to the noise stimuli was also significantly lower in the Intervention night as compared to the Disturbing night (weighted arousal score: 0.14 ± 0.04 and 0.38 ± 0.09 respectively; *t* (9) = -4.2, *p =* 0.002). Thus, despite a seemingly higher number of delivered noises in the Intervention night, the sum and probability of arousals was significantly lower in the Intervention night than in the Disturbing night.

**Figure 3.**
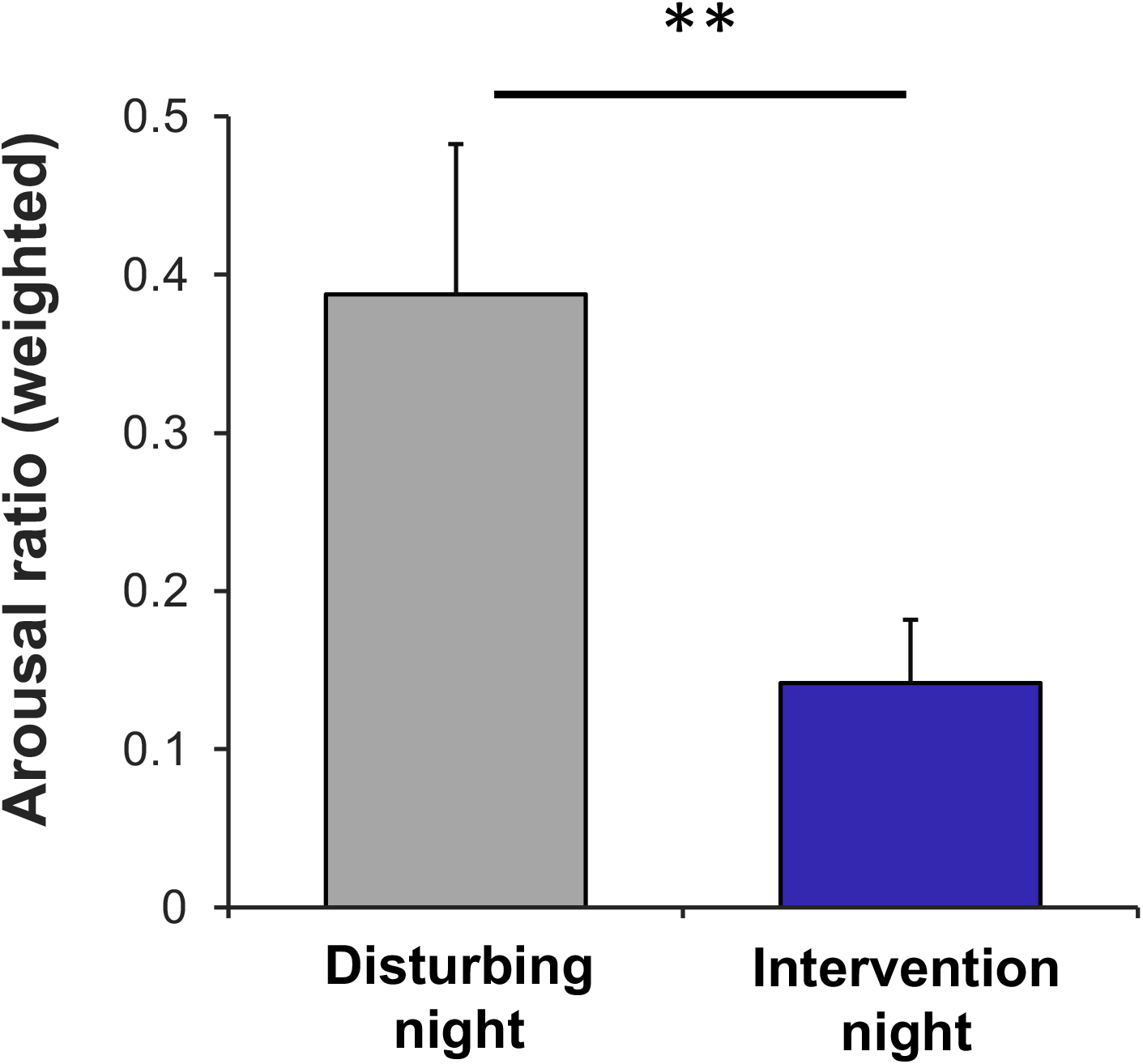
Probability of arousals due to noise. Weighted arousal scores divided by the number of delivered noises are shown (mean ± SEM; ** *p* < 0.01). The probability that an arousal would occur following a noise was strongly reduced in the Intervention night as compared to the Disturbing night. This suggests that the CLNS procedure helped protecting sleep against severely sleep-disruptive noises.

The raw number of arousals showed very similar results, which can be found as supplementary material. It is worth noting that the CLNS procedure itself never provoked any arousal.

### 3.4 Sleep architecture

Given the interactive nature of the noise presentation protocol, this study was not optimized to assess the effects of CLNS on sleep architecture. In particular, successful SO boosting due to CLNS could be immediately countered by the delivery of more noise stimuli. Despite this limitation, we did see changes in sleep architecture between the two experimental nights. Participants spent ∼10% more time in NREM in the Intervention night compared to the Disturbing night (388 ± 18 minutes and 351 ± 18 min respectively; *t* (9) =2.4, *p* = 0.03 or 70 ± 2.09 % and 65 ± 3.2 % respectively; *t* (9) = 1.54, *p* = 0.15. The duration of N1 was marginally increased in the Intervention night as compared to the Disturbing night (61 ± 17 % and 42 ± 13 % respectively; *t* (9) =2.09, *p* = 0.06). Finally, we also observed a marginal increase in Sleep efficiency in the Intervention night as compared to the Disturbing night (92 ± 5.64 % and 90 ± 6 % respectively; *t* (9) =2.07, *p* = 0.06). No other sleep architecture parameters were significantly altered (see Table 1).

**Table 1.** Sleep architecture characteristics for each experimental night.

|  | Disturbing night | Intervention night | *p* (t value) |
| --- | --- | --- | --- |
| **TST, min** | 539 ± 16 | 556 ± 17 | 0.28 (1.13) |
| **N1, min** | 42.24 ± 13 | 61.25 ± 17 | 0.06 (2.09) |
| **N1, %** | 8.7 ± 2.53 | 11 ± 3.24 | 0.08(1.19) |
| **N2, min** | 139.1 ± 14 | 147.25 ± 12.26 | 0.4 (0.87) |
| **N2, %** | 26 ± 2.75 | 26.6 ± 2.05 | 0.76 (0.31) |
| **N3, min** | 166.7 ± 24.54 | 179.7 ± 28.39 | 0.35 (0.98) |
| **N3, %** | 30.5 ± 3.91 | 33 ± 4.5 | 0.61 (0.52) |
| **NREM, min** | 351 ± 18 | 388.2 ± 18 | 0.03 (2.4) * |
| **NREM, %** | 65 ± 3.2 | 70 ± 2.09 | 0.15 (1.54) |
| **REM, min** | 188.25 ± 19.5 | 167.55 ± 11.85 | 0.28 (-1.14) |
| **REM, %** | 35 ± 3.2 | 30 ± 2.09 | 0.15 (-1.54) |
| **SE, %** | 90 ± 6 | 92 ± 5.64 | 0.06 (2.07) |
| **SL, min** | 13.4 ± 4.82 | 12 ± 3.4 | 0.19 (-1.4) |
| **REML, min** | 93.6 ± 25.89 | 85.5 ± 42.67 | 0.48 (-0.72) |
| **WASO, min** | 1.7 ± 2 | 0.5 ± 0.67 | 0.11 (-1.76) |

*Notes.* All values expressed as mean ± SEM; \* $p < 0.05$ . Percentages are calculated relative to total sleep time.
*TST* total sleep time; *SE* sleep efficiency; *REML* rapid eye movement latency; *SL* sleep latency (to stage N1); *WASO* wake after sleep onset.

### 3.5 Differences in power across frequency bands

The consideration mentioned in the previous section also applies to the power spectral analyses, such that typical CLNS induced increases in SO and spindle power may be counteracted by the interactive noise presentation protocol. Considering the whole night period, we found a significant NREM sigma power increase in the Intervention night as compared to the Disturbing night (0.026 ± 0.01 and 0.023 ± 0.008 respectively; *t* (9) = 2.3, *p* = 0.04), possibly reflecting increased spindle activity (see Figure 4A). Considering the stimulation period only, no significant differences between the two experimental nights were observed (see Figure 4B).

**Figure 4.**
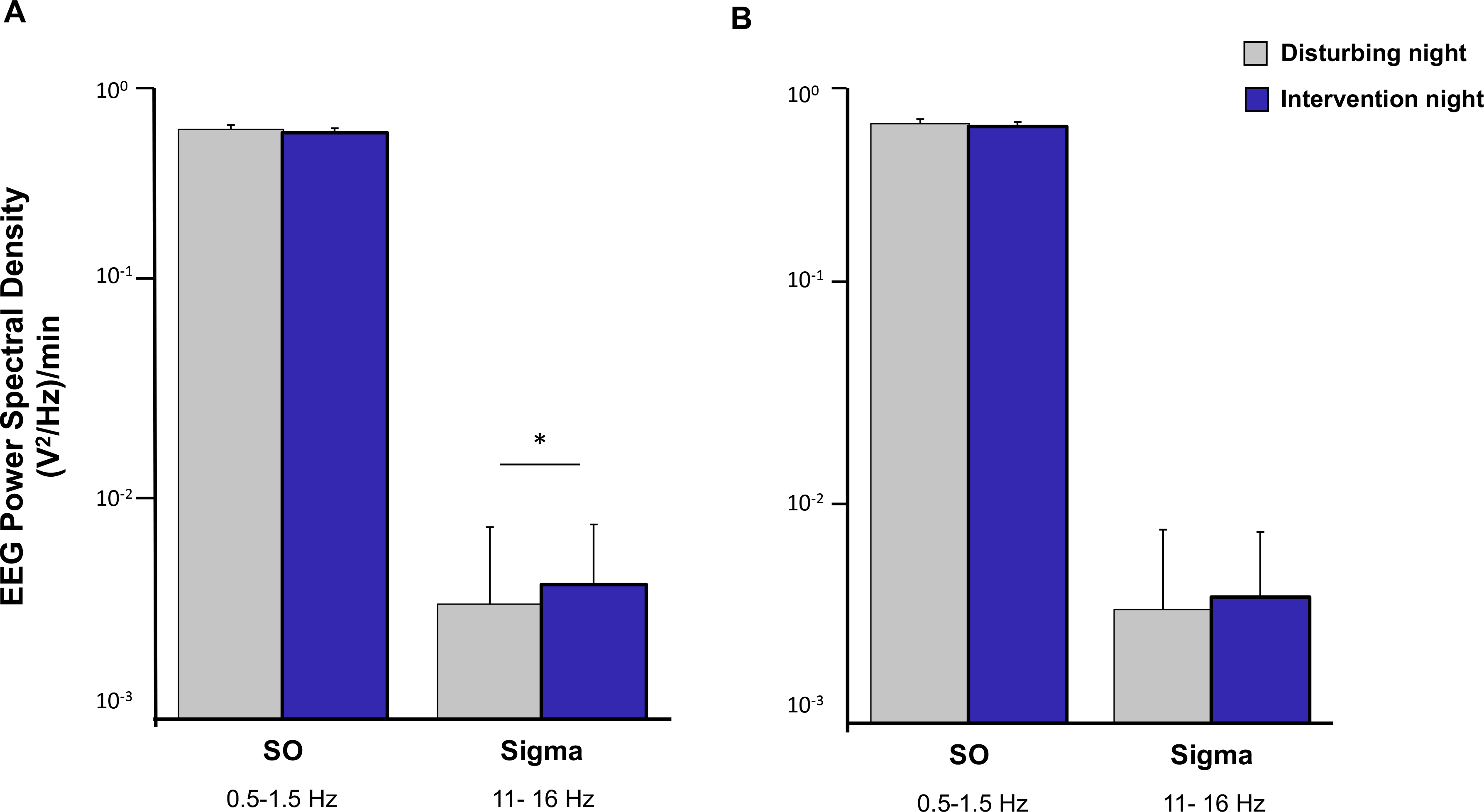
Differences in power across frequency bands in NREM sleep. (mean ± SEM relative power; * *p <* 0.05). (A) The whole night: As compared to the Disturbing nights, the power spectral density of the Intervention nights was significantly higher in the sigma band, indicative of higher sleep spindle activity. (B) The stimulation period: No differences were found for either SO or sigma bands.

### 3.6 Subjective measures of sleep

Wilcoxon’s tests on the Sleep quality questionnaire^30^, the Stanford sleepiness scale^31^ and the Consensus sleep diary^32^ revealed no significant differences between the Intervention night and the Disturbing night. Based on the questionnaires’ threshold values, participants slept equally well on both nights. Therefore, it seems that the CLNS intervention did not have any effect on subjective sleep measures (see Table 2).

**Table 2.**
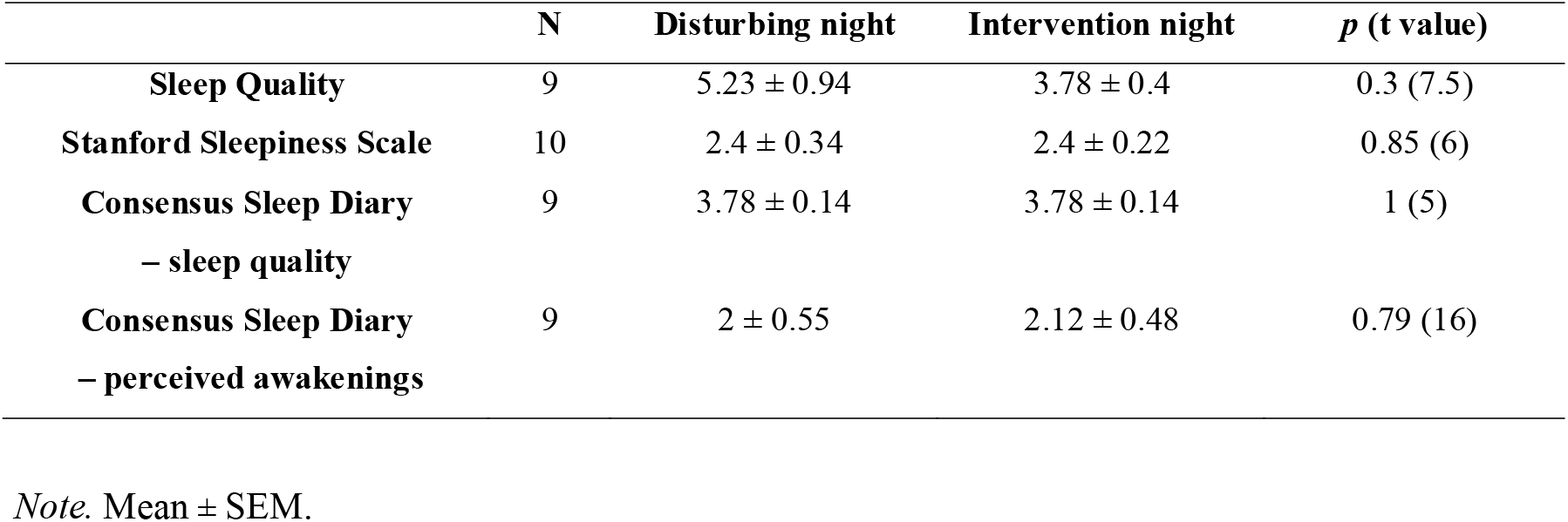
Comparison of experimental nights on subjective sleep measures.

## 4. Discussion

Exposure to noise during sleep impairs sleep quality, leading to physiological deterioration, decreased daily functioning, higher occurrence of accidents and – in the long run – increased risk for a plethora of disorders. In many circumstances it is impossible to eliminate the source of noise, leading to long-lasting distress. In this project, we tested an innovative and non-invasive approach to counteract sleep disruptions caused by noise. Taking advantage of our CLNS sleep boosting procedure, we showed that repeatedly presenting subtle sound pulses, phase-locked to 0° phase of SO during NREM sleep protects against arousals otherwise caused by environmental noise.

The sleep protective effect of the CLNS intervention was apparent in several sleep-related parameters. Most importantly, CLNS drastically decreased the occurrence of arousals caused by noise. In particular, the chance of arousals was reduced by ∼50 % in the Intervention night, compared to the Disturbing night. This provides direct evidence that subtle acoustic stimulation, phase-locked 0° phase of SO, can protect sleep continuity in the presence of noise, decoupling the sleeping brain from external acoustic stimuli.

Given the interactive nature of the noise presentation protocol, it was anticipated that successful SO boosting through CLNS would lead to the delivery of additional noise stimuli. It should be noted that this could, at least partially, counteract the finding of CLNS effects on sleep architecture and spectral content. Having said this, manipulating NREM with CLNS led to a 10% increase in time spent in NREM as compared to the Disturbing night. Additional trends for increased sleep efficiency, decreased wake time after sleep onset and increased N1, seem to support the sleep stabilizing effect of CLNS, indicating fewer full awakenings and perhaps more instances where the subject would instead remain in N1.

CLNS intervention also led to an increase in NREM sigma power compared to the Disturbing night. As sigma power is highly correlated to sleep spindle activity ^35, 36^, this might suggest an increase in spindle activity. This result points to possible neural correlates of sleep protective function of CLNS. Indeed, numerous studies support a sleep protecting role of sleep spindles^20, 37–40^. During sleep, disconnection from the external environment is maintained through a “thalamic gate”; sleep spindles are suggested to protect sleep from noise by decreasing the degree of sensory transmission through the thalamus ^39^, reducing cortical responses to sensory signals ^16^. Accordingly, in a study on healthy individuals, those with naturally high spindle density were less likely to be aroused by noise stimuli during sleep than those with low spindle density^17^. In another study, sound pulses presented during a spindle were less likely to be transmitted to higher cortical areas than sound pulses that did not coincide with a spindle ^40^. Of further interest to our results, optogenetically generating sleep spindles significantly increases the duration and stability of NREM^37^, suggesting a causal association between spindle density and NREM duration.

Of note, recent studies in our lab have shown that the CLNS procedure used here can increase delta power (1-4 Hz), as well as the amount of deep sleep in undisturbed sleepers^45^. Such sleep deepening would also be expected to stabilize sleep against disturbing influences, as thalamic inhibition and the arousal threshold are known to increase with deepening of NREM^15^ . The current study did, however, not capitalize on assessing this putative mechanism directly, as the noise delivery protocol was designed so that any enhancement of deep sleep translated directly in release of additional noises. As such, the sleep stabilizing effects of CLNS during the stimulation period are not reflected by an increase in delta power, but by a higher number of released noises.

Interestingly, a CLNS-induced increase in spindle power was observed across the whole night, but not in the stimulation period (i.e., the first two sleep cycles). This might suggest long-lasting effects of CLNS, persisting beyond the stimulation period. Possibly, early-night CLNS intervention encouraged the maintenance of stable NREM sleep for the rest of the night in an entrainment effect. Of note, previous studies evaluating effects of SO targeting on sigma power only report an evoked increase closely following the stimulus^21, 25^, and not over as long a period of unstimulated sleep as shown in this study.

As a reminder, this study used a modelling approach based on sine fitting to continuously predict the SO dynamic and deliver single acoustic stimuli targeted accurately to the 0° phase of SO. The CLNS phase targeting accuracy (phase error (mean = 3.8°, SD = 58°, VAR = 29.25°) in this study) offers very precise targeting of SO. Additionally, most previous studies delivered pairs or trains of acoustic stimuli, with only the first one targeted based on the analysis of EEG, and subsequent stimuli following the initial one at fixed 1 s intervals^43, 44^. However, phase targeting accuracy seems to be a crucial component for boosting SO and any deviation could instead lead to hinderance of SO. Indeed, presenting a second sound pulse in the down wave of the first induced SO seems to lead to a decrease in evoked SO power ^22^ and responses to rhythmic stimulation tend to quickly diminish with each subsequent sound^24^. Overall, the advantages of our method with respect to previous procedures might lead to more enduring effects.

Perhaps surprisingly, there was no difference in subjective sleep quality measures between the two nights. This could be explained by the fact that participants were able to sleep undisturbed for about 6.5 h after the 3 h of noise presentation at the beginning of the night. Thus, when recalling the quality of their sleep in the morning, participants might have only remembered the last, undisturbed part of their sleep. Given people’s notoriously poor memory for events during sleep, this is not entirely unexpected; in fact, many participants did not remember anything special about the first part of the night or had only a vague recollection of hearing noises. Also, all participants were selected to be very good sleepers, which might have steered the results towards a generalized positive appraisal of their sleep quality. Finally, as mentioned earlier, any tendency towards deeper (and thus more restorative) sleep during the stimulation period was at least partially undone by an increase in the number of noise stimuli.

Our study has some limitations. First, the small number of participants might restrict the power of the analyses. However, this was compensated by the within-subject design, which strongly limits the effects of inter-individual variance, thus guaranteeing a satisfying statistical power. In addition, included participants constituted a very homogeneous group consisting of very good, healthy sleepers in a narrow age and with similar education.

The inclusion of only very good sleepers was necessary for this proof-of-concept study to achieve a large number of arousals per participant, to enable assessment of the CLNS intervention. However, future studies should examine whether these benefits hold in other populations, including less proficient sleepers. Indeed, improving sleep in populations with sleep problems would constitute the ultimate application of the CLNS procedure. If proved effective, our CLNS intervention could eventually profit individuals beyond sleep protection, improving mental, immune and cardiovascular health^2^.

To our knowledge, this is the first experimental study to use CLNS as an application strategy to stabilize sleep in the presence of noise. Our results show that CLNS intervention strongly reduces the number of arousals to noise, possibly through a mechanism involving increased spindle activity and sensory decoupling.

The results from this study show that CLNS can be used to precisely phase-lock non-arousing sound pulses to the 0° phase of SO, even in the presence of noise-induced sleep disturbance, possibly through a mechanism involving increased spindle activity and sensory decoupling. Importantly, this intervention can successfully protect sleep from disruption caused by environmental noise. CLNS is a safe, non-invasive procedure that is currently being tested for applicability in naive users at home, using a portable tool. As such, CLNS has the potential to support recovery in hospitalized patients, improve daily functioning in high-demanding jobs and bears crucial relevance for areas where noise cannot be easily lowered to an acceptable level.

## Acknowledgements

We thank Dominik Koller and Ilia Korjourov for their help with aspects of the study design.

## References

1. Irwin MR. Why Sleep Is Important for Health: A Psychoneuroimmunology Perspective. Annu Rev Psychol. 2015;66(1):143–172. doi:10.1146/annurev-psych-010213-115205

2. Grimaldi D, Papalambros NA, Reid KJ, et al. Strengthening sleep-autonomic interaction via acoustic enhancement of slow oscillations. Sleep. 2019;42(5). doi:10.1093/sleep/zsz036

3. Van Dongen HPA, Maislin G, Mullington JM, Dinges DF. The cumulative cost of additional wakefulness: dose-response effects on neurobehavioral functions and sleep physiology from chronic sleep restriction and total sleep deprivation. Sleep. 2003;26(2):117–126. doi:10.1093/sleep/26.2.117

4. Luyster FS, Strollo PJJ, Zee PC, Walsh JK. Sleep: a health imperative. Sleep. 2012;35(6):727–734. doi:10.5665/sleep.1846

5. Weinger MB, Ancoli-Israel S. Sleep Deprivation and Clinical Performance. JAMA. 2002;287(8):955-957. doi:10.1001/jama.287.8.955

6. Lockley SW, Barger LK, Ayas NT, Rothschild JM, Czeisler CA, Landrigan CP. Effects of Health Care Provider Work Hours and Sleep Deprivation on Safety and Performance. Jt Comm J Qual Patient Saf. 2007;33(11, Supplement):7-18. doi:https://doi.org/10.1016/S1553-7250(07)33109-7

7. Chaput J-P, Dutil C, Sampasa-Kanyinga H. Sleeping hours: what is the ideal number and how does age impact this? Nat Sci Sleep. 2018;10:421–430. doi:10.2147/NSS.S163071

8. Hume K, Brink M, Basner M. Effects of environmental noise on sleep YR - 2012/12/1. Noise Heal. (61 UL-https://www.noiseandhealth.org/article.asp?issn=1463-1741;year=2012;volume=14;issue=61;spage=297;epage=302;aulast=Hu;t=5):297 OP-302 VO - 14. doi:10.4103/1463-1741.104897

9. Buxton OM, Ellenbogen JM, Wang W, et al. Sleep Disruption due to Hospital Noises. Ann Intern Med. 2012;157(3):170–179. doi:10.7326/0003-4819-156-12-201208070-00472

10. Jones K. Aircraft noise and sleep disturbance: A review. 2009:30.

11. Alsulami G, Rice AM, Kidd L. Prospective repeated assessment of self-reported sleep quality and sleep disruptive factors in the intensive care unit: acceptability of daily assessment of sleep quality. BMJ Open. 2019;9(6):e029957. doi:10.1136/bmjopen-2019-029957

12. Barger L, Flynn-Evans E, A WK, et al. Prevalence of Sleep Deficiency and Hypnotic Use Among Astronauts Before, During and After Spaceflight: An Observational Study. Aviat Space Environ Med. 2014;13(9):904–912. doi:10.1016/S1474-4422(14)70122-X.Prevalence

13. Dijk DJ, Neri DF, Wyatt JK, et al. Sleep, performance, circadian rhythms, and light-dark cycles during two space shuttle flights. Am J Physiol Regul Integr Comp Physiol. 2001;281(5):R1647–64. doi:10.1152/ajpregu.2001.281.5.R1647

14. Wu B, Wang Y, Wu X, Liu D, Xu D, Wang F. On-orbit sleep problems of astronauts and countermeasures. Mil Med Res. 2018;5(1):1–12. doi:10.1186/s40779-018-0165-6

15. Neckelmann D, Ursin R. Sleep stages and EEG power spectrum in relation to acoustical stimulus arousal threshold in the rat. Sleep. 1993;16(5):467–477.

16. Steriade M. Neuronal Substrates of Sleep and Epilepsy. Cambridge University Press; 2003.

17. Dang-Vu TT, McKinney SM, Buxton OM, Solet JM, Ellenbogen JM. Spontaneous brain rhythms predict sleep stability in the face of noise. Curr Biol. 2010;20(15):626–627. doi:10.1016/j.cub.2010.06.032

18. Perrault AA, Khani A, Quairiaux C, et al. Whole-Night Continuous Rocking Entrains Spontaneous Neural Oscillations with Benefits for Sleep and Memory. Curr Biol. 2019;29(3):402–411.e3. doi:10.1016/j.cub.2018.12.028

19. Marshall L, Helgadóttir H, Mölle M, Born J. Boosting slow oscillations during sleep potentiates memory. Nature. 2006;444(7119):610–613. doi:10.1038/nature05278

20. Lustenberger C, Boyle MR, Alagapan S, Mellin JM, Vaughn B V., Fröhlich F. Feedback-Controlled Transcranial Alternating Current Stimulation Reveals a Functional Role of Sleep Spindles in Motor Memory Consolidation. Curr Biol. 2016;26(16):2127–2136. doi:10.1016/j.cub.2016.06.044

21. Besedovsky L, Ngo H-V V, Dimitrov S, Gassenmaier C, Lehmann R, Born J. Auditory closed-loop stimulation of EEG slow oscillations strengthens sleep and signs of its immune-supportive function. Nat Commun. 2017;8(1):1984. doi:10.1038/s41467-017-02170-3

22. Cox R, Korjoukov I, De Boer M, Talamini LM. Sound asleep: Processing and retention of slow oscillation phase-targeted stimuli. PLoS One. 2014;9(7). doi:10.1371/journal.pone.0101567

23. Ngo HV V., Claussen JC, Born J, Mölle M. Induction of slow oscillations by rhythmic acoustic stimulation. J Sleep Res. 2013;22(1):22–31. doi:10.1111/j.1365-2869.2012.01039.x

24. Ngo HV V., Martinetz T, Born J, Mölle M. Auditory closed-loop stimulation of the sleep slow oscillation enhances memory. Neuron. 2013;78(3):545–553. doi:10.1016/j.neuron.2013.03.006

25. Ngo H-V V., Miedema A, Faude I, Martinetz T, Molle M, Born J. Driving Sleep Slow Oscillations by Auditory Closed-Loop Stimulation--A Self-Limiting Process. J Neurosci. 2015;35(17):6630–6638. doi:10.1523/JNEUROSCI.3133-14.2015

26. Ong JL, Lo JC, Chee NIYN, et al. Effects of phase-locked acoustic stimulation during a nap on EEG spectra and declarative memory consolidation. Sleep Med. 2016;20:88–97. doi:10.1016/j.sleep.2015.10.016

27. Papalambros NA, Santostasi G, Malkani RG, et al. Acoustic Enhancement of Sleep Slow Oscillations and Concomitant Memory Improvement in Older Adults. Front Hum Neurosci. 2017;11(March):1–14. doi:10.3389/fnhum.2017.00109

28. Talamini LM, Juan E. Sleep as a window to treat affective disorders. Curr Opin Behav Sci. 2020;33(Figure 2):99–108. doi:10.1016/j.cobeha.2020.02.002

29. Academy of Sleep Medicine A. The AASM Manual for the Scoring of Sleep and Associated Events Summary of Updates in Version 2.5. J Clin Sleep Med. 2020:1–2.

30. Simor P, Köteles F, Bodizs R, Bardos G. A questionnaire based study of subjective sleep quality: The psychometric evaluation of the Hungarian version of the Groningen Sleep Quality Scale. Mentálhigiéné és Pszichoszomatika. 2009;10. doi:10.1556/Mental.10.2009.3.5

31. Shahid A, Wilkinson K, Marcu S, Shapiro CM. Stanford Sleepiness Scale (SSS) BT - STOP, THAT and One Hundred Other Sleep Scales. In: Shahid A, Wilkinson K, Marcu S, Shapiro CM, eds. New York, NY: Springer New York; 2012:369–370. doi:10.1007/978-1-4419-9893-4_91

32. Carney CE, Buysse DJ, Ancoli-Israel S, et al. The consensus sleep diary: standardizing prospective sleep self-monitoring. Sleep. 2012;35(2):287–302. doi:10.5665/sleep.1642

33. Berry RB, Brooks R, Gamaldo C, et al. AASM scoring manual updates for 2017 (version 2.4). J Clin Sleep Med. 2017;13(5):665–666. doi:10.5664/jcsm.6576

34. Planck M, Berens P, Velasco MJ. The circular statistics toolbox for Matlab The circular statistics toolbox for Matlab. Biol Cybern. 2009;(184).

35. Fearon RMP, Reiss D, Leve LD, et al. HHS Public ccess. Dev Psychopathol. 2015;27(4):1251-1265. doi:10.1038/nmeth.2855.Sleep

36. Warby SC, Wendt SL, Welinder P, et al. Sleep-spindle detection: crowdsourcing and evaluating performance of experts, non-experts and automated methods. Nat Methods. 2014;11(4):385–392. doi:10.1038/nmeth.2855

37. Kim A, Latchoumane C, Lee S, et al. Optogenetically induced sleep spindle rhythms alter sleep architectures in mice. Proc Natl Acad Sci U S A. 2012;109(50):20673–20678. doi:10.1073/pnas.1217897109

38. Wimmer RD, Astori S, Bond CT, et al. Sustaining sleep spindles through enhanced SK2-channel activity consolidates sleep and elevates arousal threshold. J Neurosci. 2012;32(40):13917–13928. doi:10.1523/JNEUROSCI.2313-12.2012

39. Elton M, Winter O, Heslenfeld D, Loewy D, Campbell K, Kok A. Event-related potentials to tones in the absence and presence of sleep spindles. J Sleep Res. 1997;6(2):78–83. doi:10.1046/j.1365-2869.1997.00033.x

40. Dang-Vu TT, Bonjean M, Schabus M, et al. Interplay between spontaneous and induced brain activity during human non-rapid eye movement sleep. Proc Natl Acad Sci U S A. 2011;108(37):15438–15443. doi:10.1073/pnas.1112503108

41. Göldi M, van Poppel EAM, Rasch B, Schreiner T. Increased neuronal signatures of targeted memory reactivation during slow-wave up states. Sci Rep. 2019;9(1):1–10. doi:10.1038/s41598-019-39178-2

42. Krugliakova E, Volk C, Jaramillo V, Sousouri G, Huber R. Changes in cross-frequency coupling following closed-loop auditory stimulation in non-rapid eye movement sleep. Sci Rep. 2020;10(1):10628. doi:10.1038/s41598-020-67392-w

43. Bellesi M, Riedner BA, Garcia-Molina GN, Cirelli C, Tononi G. Enhancement of sleep slow waves: Underlying mechanisms and practical consequences. Front Syst Neurosci. 2014;8(October):1–17. doi:10.3389/fnsys.2014.00208

44. Garcia-Molina G, Tsoneva T, Neff A, et al. Hybrid in-phase and continuous auditory stimulation significantly enhances slow wave activity during sleep. Proc Annu Int Conf IEEE Eng Med Biol Soc EMBS. 2019:4052–4055. doi:10.1109/EMBC.2019.8857678

45. Talamini, T. (2019). Enhancing or depressing memories, while deepening sleep, by EEG-guided neurostimulation. World Sleep 2019, Vancouver, Canada (paper in preparation).

