## Supplementary table 1 for "The looping lullaby: closed-loop neurostimulation decreases sleepers’ sensitivity to environmental noise"

**Supplementary material**

|  | Arousal ratio (raw) |
| --- | --- |
| 1 | 0.309499 |
| 2 | 0.134262 |

**Supplement: Figure 1. Probability of raw arousals in the stimulation period** (mean ± SEM; ** *p* < 0.01). The number of arousals was reduced by ~52% in the Intervention night as compared to the Disturbing night (4.7 ± 0.83 and 9 ± 1.25 respectively; *t* (9) = -2.61, *p* = 0.02) (Supplementary Figure 1). This result was further supported with a decrease in the probability of arousals due to noise in the Intervention night as compared to the Disturbing night (0.13 ± 0.03 and 0.3 ± 0.05 respectively; *t* (9) = -6.28, *p* <.001).
